# Comparative physiology of five tropical montane songbirds reveals differential seasonal acclimatisation and cold adaption

**DOI:** 10.1101/2020.05.22.111328

**Authors:** Samuel E.I. Jones, Martin Suanjak, Joseph A. Tobias, Robin Freeman, Steven J. Portugal

**Affiliations:** Department of Biological Sciences, School of Life and Environmental Sciences, Royal Holloway University of London, Egham, Surrey TW20 0EX; Institute of Zoology, Zoological Society of London, Regents Park, London NW1 4RY; Department of Tropical Ecology and Animal Biodiversity, University of Vienna, Universitätsring 1, 1010 Wien, Vienna, Austria; Department of Life Sciences, Imperial College London, Silwood Park, Buckhurst Road, Ascot, Berkshire SL5 7PY

**Keywords:** avian metabolism, elevational range, pace of life, phenotypic flexibility, thermoregulation

## Abstract

The physiology of tropical birds is poorly understood, particularly in how it relates to local climate and changes between seasons. This is particularly true of tropical montane species, which may have sensitive thermal tolerances to local microclimates. We studied metabolic rates (using open flow respirometry), body mass and haemoglobin concentrations of five sedentary Mesoamerican songbirds between the summer and winter at two elevations (1550 m and 1950 m, respectively). We asked whether there were uniform seasonal shifts in physiological traits across species, and whether higher elevation species displayed evidence for cold tolerance. Seasonal shifts in metabolic rates differed between the three species for which data were collected. Basal metabolic rates in one species – black-headed nightingale-thrushes *Catharus mexicanus* – were up-regulated in summer (~19% increase of winter metabolism), however two other species displayed no seasonal regulation. No species exhibited shifts in haemoglobin concentrations across season or across elevation, whereas body mass in two species was significantly higher in the summer. One species restricted to higher elevations – ruddy-capped nightingale-thrushes *C. frantzii* – displayed physiological traits indicative of cold-tolerance. Although only summer data were available for this species (*C. frantzii*), metabolic rates were constant across temperatures tested (5-34°C) and haemoglobin concentrations were significantly higher compared to the other four species. Our results suggest that seasonal acclimatisation in physiological traits is variable between species and appear unrelated to changes in local climate. As such, the distinct physiological traits observed in ruddy-capped nightingale-thrushes likely relate to historic isolation and conserved physiological traits rather than contemporary climatic adaption.

## Introduction

Physiological acclimatisation, where the physiological characteristics of a species are shaped by local climate, is intrinsic to an organisms ability to survive in changeable environments (Chown *et al.* 2004, Bozinovic *et al.* 2011). In birds, non-migratory species are frequently used to investigate such physiological shifts, due to the temperature extremes they typically experience throughout the duration of the annual cycle (McKechnie 2007, Swanson 2010). Consistent physiological acclimatisation to cold winter temperatures have been displayed in resident high-latitude birds, such as winter increases in basal and peak metabolic rates (BMR/PMR; the lower and upper limits of metabolic power output, respectively) and body mass (Mb), reflecting the increased energetic demands of maintaining high internal body temperatures (McKechnie 2007, McKechnie & Swanson 2010, Smit & McKechnie 2010, McKechnie *et al.* 2015). However, while the general patterns of physiological acclimatisation to cooler climates are relatively well established in temperate birds (Swanson 2010), much less is known in lower latitude species.

How avian physiology relates to the environmental conditions in tropical latitudes is poorly understood, and is largely restricted to studies in warm tropical lowlands (Pollock *et al.* 2019). Tropical birds have lower metabolic rates compared to those of temperate species (Wiersma *et al.* 2007, Londoño *et al.* 2015, Bushuev *et al.* 2017), suggested to be an ecophysiological adaption to the warmer and more seasonally stable climates across the tropics, in contrast to the cooler and more seasonally variable climates at temperate latitudes (White *et al.* 2007, Jetz *et al.* 2008, Khaliq *et al.* 2014). For example, tropical birds typically have narrower thermo-neutral zones (TNZ; defined by the upper and lower temperature bounds at which a species begins thermoregulation) than temperate species, reflecting reduced demands of thermogenesis in less seasonal and warmer climates (e.g. Khaliq *et al.* 2015). Whether the physiology of tropical birds is actually a product of reduced climatic variability has subsequently been questioned, however, where the BMR of over 250 species in the Peruvian Andes did not differ across elevation despite the cooler and more variable environmental conditions at higher elevations (Londoño *et al.* 2015, 2017).

Two studies investigating seasonal variation in metabolic rates have also indicated that the physiology of tropical birds may be unrelated to environmental temperatures (McKechnie *et al.* 2015, Pollock *et al.* 2019). Both Pollock et al. (2019) and Wells & Schaeffer (2012) found considerable variation in BMR, PMR, and thermoregulatory traits in lowland Panamanian rainforest species between the summer and winter months. The direction of the variability in these studies was not consistent between species, however, with some taxon displaying no seasonal acclimatisation in BMR or thermoregulatory traits, while others increased or decreased metabolic rates in the winter, despite the lack of seasonal temperature fluctuations (Pollock *et al.* 2019). Similarly, Wells & Schaeffer (2012) found decreased PMR in tropical birds during the winter months, the opposite of which is generally found in temperate species. Taken together, these studies suggest that the ecophysiology of tropical species is dictated by factors other than temperature (McKechnie *et al*. 2015). The ubiquity of this hypothesis remains unclear, however, because comparative studies are lacking away from tropical lowland forests where climatic variation is generally low.

Tropical mountains offer valuable case studies in avian physiology because they allow assessments of whether physiological traits of resident species relate to variable climates (Londoño *et al.* 2015). Changes in elevation along tropical mountains are characterised by sharp changes in temperature isotherms and it has been historically hypothesised that tropical montane species have evolved distinct physiological tolerances unique to their elevational distributions (Janzen 1967, Ghalambor *et al*. 2006). Thus, if tropical montane species are physiologically sensitive, the physiological traits of sedentary species may also reflect changes in seasonal conditions. To explore this further, we assessed whether cooler environmental temperatures at higher elevations in tropical mountains manifest in physiological cold-tolerance traits in resident species, and whether species displayed uniform shifts between seasons.

We compared metabolic rates (both BMR and thermoregulation in response to manipulated temperatures), M_b_, and total blood haemoglobin concentrations (H_b_) of five sedentary highland Central American songbirds. The BMR of a species is a widely used measure in avian physiology, reflecting the lowest energetic requirements for homeostasis, while thermoregulation reflects the metabolic responses to changing temperatures (Lighton 2008). In addition, M_b_ and H_b_ reflect fluctuations in physiological condition related to both oxidative stress (H_b_) and cold tolerance (Swanson 2010, Labocha & Hayes 2011, Minias 2015, 2020), allowing us to assess whether any observed changes in these traits also related to changes in condition. We asked whether our study species responded to cooler winter conditions (non-breeding season) by increasing BMR, M_b_ and H_b_, and whether these changes were consistent between species. Additionally, we asked whether a higher elevation species displayed physiological differences attributable to colder conditions at higher elevations such as higher H_b_ content, increased BMR and lower temperature limits to the TNZ.

## Methods and Materials

### Study site and focal species

We measured BMR, M_b_, H_b_ and thermoregulation in black-headed nightingale-thrushes *Catharus mexicanus,* ruddy-capped nightingale-thrushes *C. frantzii,* chestnut-capped brushfinches *Arremon brunneinucha,* grey-breasted wood wrens *Henicorhina leucophrys,* and H_b_ and M_b_ only in common bush tanagers *Chlorospingus flavopectus* (hereafter, BHNT, RCNT, CCBF, GBWW and COBT, respectively) in Cusuco National Park, in the Sierra del Merendón, north-western Honduras (approximately N15.552, E – 88.296). Fieldwork was undertaken between June-August, 2017/18 and January 2018.

The study species breed during the warmer months of May-August (summer) at the field site (see Howell & Webb 1995), while January (winter) is cooler. We measured environmental temperature (T_e_; °C) at two research camps (1550m and 1950m, respectively-see below) using remote loggers (HOBO UA-001-64, Onset, USA, data pooled from 2-3 loggers per camp) deployed for the entirety of the study period in order to capture seasonal and elevational T_e_ variation. Loggers were attached to trees ~1m off the forest floor to best represent the T_e_ experienced by the study species. Mean T_e_ (°C ± SD) during the summer at 1550m was 17.4 ± 1.7 (max = 24.7, min = 12.2), and at 1950m was 15.5 ± 1.4 (max = 22, min = 11.2). During the winter, T_e_, at 1550m was 13.2 ± 2.2 (max = 21.2, min = 7.8), and at 1950m was 11.3 ± 2.11 (max = 16.3, min = 5.24) (Fig. S1).

We studied birds at both research camps, but because of logistical difficulties, all winter fieldwork was undertaken at the lower elevation (1550m) camp only. Each species in our study occurs at both camps except RCNT (which occurs only at higher elevations at the study site – see Jones *et al.* (2019)). Accordingly, we only obtained summer data on RCNT. None of the study species undertake any known seasonal elevational movements at the study site (determined by both consistent captures/re-sightings of banded birds at their site of original capture, and/or year-round territory occupation throughout the course of the study).

### Capture and handling

We captured birds using 6 or 9 m mist-nets, generally lured into the nets by conspecific playback. This method inherently targeted territorial holders and as such our sample is biased towards male birds. After capture, each bird was measured for a standard set of biometrics (e.g. maximum wing chord and tarsus length) and M_b_ (in g) to an accuracy of 0.1g using digital scales (SA-500, SATRUE, Taiwan). Each bird was banded with a uniquely numbered aluminium ring (Aranea, Łódź, Poland) and unique field-identifiable combination of coloured rings for future study.

Birds were aged and sexed, where possible, by a combination of plumage dimorphism and moult limits, and checked for evidence of breeding condition (enlarged cloacal protuberance and brood patches). During the summer, most males of the study species (particularly BHNT, RCNT, CCBF) were in reproductive condition (enlarged cloacal protuberances). Breeding seasons for the study species are prolonged, and as such, it is generally not possible to sample birds in the summer season not in some state of reproductive condition. We did not include juvenile birds in any of our samples, although we did include first-cycle birds (i.e. born the previous breeding season), because they are functionally adult (e.g. Wolfe *et al.* 2010). Fat and muscle content were also assessed, but variance was virtually indistinguishable between season (where fat reserves were rarely present and muscle profiles were constant). These data were thus not suitable for statistical tests. Female birds with edematous brood patches were released after processing for ethical reasons. Birds that were transferred to the respirometry system (see below) for metabolic measurements were caught between 15:00 and 18:30 (dusk).

### Blood sampling

We measured H_b_ concentration in grams per decilitre (g/dL) using a portable analyser (Prospect haemoglobin, prospect Diagnostics Ltd, UK). H_b_ concentrations reflect the ability of a bird to meet its oxygen requirements and as such it is a measure of physiological condition and cold adaption for both between and within species comparisons (Dubay & Witt 2014, Minias 2015). Blood was sampled shortly after capture from the alar vein and drained directly into a reagent-free cuvette (<8uL) specific to the analyser. After blood was drained into the cuvette, pressure was applied to the vein using cotton wool to stem the bleeding (see procedures described by Owen (2011)). Each blood sample was tested three times in the unit, with the resulting values averaged. The accuracy of the H_b_ analyser was measured every 14 days throughout fieldwork using standardised control measures (DiaSpect Control, DiaSpect Mecidal, Germany) at three concentrations: 8.0±0.4 g/dL, 12.6±0.6 g/dL and 16.0±0.8 g/dL.

### Respirometry

We measured energy metabolism (as a rate of oxygen consumption 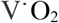 ml/min^1^) using open flow respirometry with a portable gas analysis system (FoxBox, Sable Systems, USA; hereafter ‘FoxBox’). Birds were placed in a custom-built 20cm^3^ Perspex chamber fitted with a perch into which a constant airflow was supplied. We experimentally manipulated temperatures by placing the respirometry chamber inside a modified cooler box fitted with a Peltier thermoelectric cooler module (AC-046, TE Technology, USA), capable of heating/cooling the interior of the cool box by a temperature controller (TC-48-20, TE Technology, USA). We checked temperatures inside the chamber with a logger (HOBO UA-001-64, Onset, USA) and offset slight differences between the temperature set inside the cooler box and inside the chamber using the relevant setting on the temperature controller. Temperature plates were powered by an analogue 24V power bench (ALF2412, ELC, UK), externally powered by a portable generator.

Ambient air was dried (self-indicating silica gel, GeeJay chemicals, UK) and pulled through the chamber at 1000±1 ml/min; a sufficient flow rate at which O_2_ levels did not fall below 0.5% of natural levels, preventing hypoxia/hypercapnia. Air flow was measured and controlled by a linearized mass flow meter internal to the FoxBox, meaning temperature and barometric pressure compensation to STP were not required. Excurrent air from the chamber was dried again (silica gel) before entering the FoxBox where O_2_ and CO_2_ content were recorded at 1s intervals (Sable Systems ExpeData, Las Vegas, USA). A second air channel (outside ambient air, taken directly adjacent to incurrent air to chamber) for use as reference ambient air was manually routed to the FoxBox following each temperature treatment (see below) to correct for drift in the analysers. Before sessions began, the O_2_ analyser was spanned to 20.95% (environmental O_2_ concentration) after gas measurements had stabilised.

### Experimental procedure and temperature manipulations

All respirometry sessions were undertaken at night to ensure birds were in their natural resting circadian phase. Following capture, we roosted all birds in cloth bags in a quiet room where they were fasted (without food). Birds were transferred to the respirometry system after dusk (1845 onwards) and left to acclimatise to the chamber for at least 45mins before temperature manipulations began. This acclimatisation period was sufficient washout time for residual chamber air to be replaced (see Lighton 2008).

Each bird was subjected to 1-4 temperature treatments per night. Temperature treatments for each bird were randomly chosen within five bands: 5-10°C, 11-16°C, 17-22°C, 23-28°C, and 29-34°C, the treatment temperatures within which were systematically rotated (e.g. within 5-10°C, 5°C, then 6°C etc. to avoid repeated measures at the same temperatures). We did not exceed 34°C during temperature manipulations as the focus was cold tolerance in the study species (the lower temperature limit of thermal-neutrality), and 34°C was likely to be within thermal-neutrality for our study species (Londoño *et al.* 2017). Once each temperature treatment was stable (±0.5°C of target temperature), the bird was left for a further 45 minutes at each temperature before data were accepted for metabolic measurements. We then recorded data for ~20 minutes before setting the next temperature treatment. Baselines of 7 minutes of ambient air were taken before each new temperature treatment was set (exact time intervals varied but were typically every 1-1.5hrs). All metabolic data throughout the study were taken at least four hours after capture to ensure birds were post-absorptive (Karasov 1990). Respirometry sessions typically finished by 03:00, and birds were released at the site of capture the following morning.

### Data processing

Respirometry traces were baseline corrected and converted to 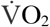 (ml/min^-1^) using equation 11.7 in Lighton (2008) (equation (1) below):

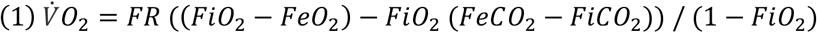

Where, FR= Flow Rate and FiO_2_/CO_2_/FeO_2_/CO_2_= incurrent and excurrent fractional concentrations of O_2_, and CO_2_, respectively. Incurrent Oxygen (FiO_2_) is at atmospheric levels (0.2095%). 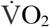 at each temperature trial was taken as the lowest continuous average over three minutes when the trace was low and stable (all values ≤0.05% of the mean), and where temperature had been constant for 45-minutes (see above). We subsequently converted 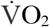 to Watts (W) using a joule conversion of 20.1 J mL^-1^ (Gessaman & Nagy 1988).

### Statistical analysis

All analyses where conducted in ‘R’ (R Core Team 2016). Values are presented as mean ± standard deviation (SD) or standard error (SEM) for model parameters, and statistically significance thresholds as *p* values <0.05. We used a mixed modelling approach for all analyses due to the repeated measures either inherent to the experimental design (e.g. metabolic measurements of individuals at multiple treatment temperatures), or because of the repeated measures on individuals between seasons.

We analysed metabolic rates in a two-step process. Firstly we fitted linear mixed effects models for each species using the lme4 package (Bates *et al.* 2014). We modelled metabolic rate (W) as a function of season (summer/winter), M_b_ (g), experimental temperature treatment (°C) and elevation (m) (of the captured bird) and a season × experimental temperature interaction as fixed effects, and included bird identity (individual) as a random effect. The factor ‘season’ was not included for RCNT models because no winter data were collected for this species. In incidences when M_b_ was identified as a significant effect in full models (BHNT and CCBF), we repeated the analyses using mass-specific metabolic rate as the response variable. All continuous explanatory variables were scaled to account for potential sensitivity in magnitudes of change within variables. We then selected best fitting models for each species by dredging all possible model iterations, ranking the resulting models by corrected Akaike Information Criterion (AICc) in the MuMIn package (Barton 2016). We then examined models for significant parameters with Wald-Chi square tests using the *car* package (Fox *et al*. 2011). All models within 6Δ AICc of the best-fitting model, as well as the saturated and null models per species are presented in the supplementary materials. Hereafter, ‘best-fitting model’ refers to that with 0Δ AICc, while ‘top model set’ refers to all models within 6Δ AICc. When the best-fitting model had multiple parameters, parsimonious models for each significant effect in this model are also presented. This modelling approach broadly follows recommendations suggested by Harrison *et al.* (2018).

Where treatment temperature was identified as a significant parameter in the top model set, we then fitted non-linear mixed models to the data in the nlme package (Pinheiro *et al*. 2017). These models estimated how metabolic rates were affected by temperature by estimating the inflection temperature (lower critical limit of thermoneutrality – T_1c_) at which species began thermoregulation (i.e. an increase in metabolic rate), the slope of this relationship (minimum thermal conductance – C_min_) and metabolic rate above T_1c_ (BMR). This method can underestimate minimum thermal conductance (see McNab 1980), but because we did not measure body temperatures, this allowed an approximation of energetic costs of thermoregulation below T_1c_. Because of this, we elected not to statistically compare metabolic responses to temperature, instead qualitatively comparing values of C_min_ and T_1c_ between species. In one species (BHNT), where seasonal differences in metabolic rates where apparent from model selections, we added a two-way factor of ‘season’ (summer/winter) in order to assess specifically which parameters (BMR, C_min_ or T_1c_) differed between season.

We analysed H_b_ using the same modelling process as described for the first step of metabolic data. We predicted H_b_ (g/dL) as a function of sex (male/female), season (summer/winter), elevation (m) and a sex × season interaction as fixed effects, with bird identity (individual) as a random effect. The factors ‘sex’ and ‘season’ for RCNT, and ‘sex’ for GBWW were not included for these models, as these data were not available. After assessing intraspecific differences in H_b_ per species, we assessed interspecific differences with post-hoc comparison tests using the multcomp package (Bretz *et al*. 2010) on a simpler mixed model of H_b_ as a function of species. We compared summer and winter datasets separately, however, to compare like-for-like data. Finally, we tested for seasonal differences in M_b_ using Wald-Chi square tests on linear mixed models of M_b_ as a function of season. For BHNT – the only species we were able to discern sex between season – we also tested for sex specific season changes in M_b_.

To place our results within the broader context of the physiological diversity of tropical birds, we compared BMR of the species in our study to predicted values from mass-scaling exponents, using phylogenetically informed power equations in Londoño *et al.* (2015). Values are presented as percentages of predicted BMR, using coefficients presented for tropical species when those with ambiguous breeding distributions were excluded from the dataset (see Londoño *et al.* 2015). For species in our study that displayed no seasonal changes in BMR, we pooled data across seasons. We considered values within 10% of those predicted as within the expected range broadly following similar studies (e.g. Smit & McKechnie 2010). We estimated BMR for each species in our study by taking the lowest measure of metabolic rate per individual above T_1c_, comparable to similar studies (Londoño *et al.* 2015). Finally, for purposes of comparison, we present summer and winter values for BMR (both whole animal and mass-specific), H_b_ concentrations, and M_b_ and calculate winter/summer ratios by dividing mean winter values by mean summer values (McKechnie 2007, Pollock *et al.* 2019).

## Results

### Metabolism; responses to temperature and seasonal change

We found a strong negative effect of temperature on metabolic rates for three species in the best fitting models (BHNT χ^2^_1_ = 58.708, *P* <0.001; CCBF χ^2^_1_ = 29.19, *P* <0.001; GBWW χ^2^_1_ = 19.72, *P* <0.001) but not RCNT (Fig. 1). Metabolic rate (W) was best predicted by temperature alone for CCBF and GBWW, for BHNT, however, season was also a significant covariate (χ^2^_1_ = 21.872, *P* <0.001) in addition to temperature. For RCNT, none of the variables tested influenced metabolic rates and the best-fitting model was one with the intercept alone. We found elevation had no effect on metabolic rate for any of the study species, although for RCNT elevation was significant in the second ranked model (χ^2^_1_ = 4.056, *P* = 0.044). M_b_ was a significant variable in full and top model sets for both BHNT and CCBF, but when the same analyses were undertaken on mass-corrected values for these species, our results were comparable. Because of this, we fitted subsequent models on for these two species on whole-animal values. For full model selection results for each species (including mass corrected fits), see Tables S1-4.

**Fig.**
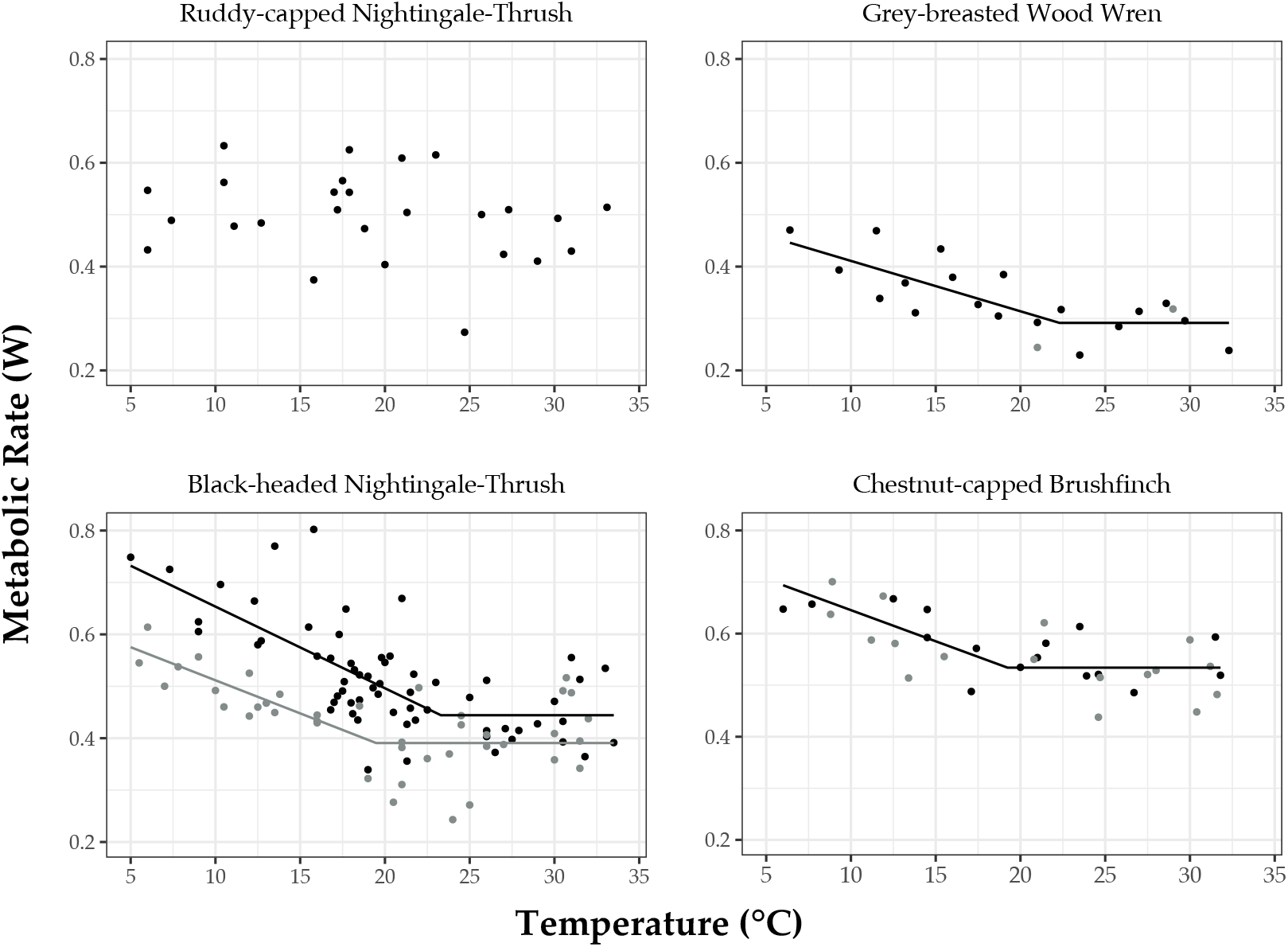
Error! No text of specified style in document.. Metabolic rate (W) as a function of temperature (°C) for four species of cloud-forest songbirds. Data points are temperature trials and fitted lines are non-linear mixed model fits estimating the inflection temperature at which thermoregulation began (T_1c_), minimum thermal conductance (C_min_) and metabolic rates above the inflection temperature (BMR). Summer (black) and winter (grey) data are plotted separately, but summer and winter model fits are separated for black-headed nightingalethrushes *Catharus mexicanus*. No effect of temperature was apparent for ruddy-capped nightingale-thrushes *Catharus frantzii* (top left panel).

Lower critical temperatures (T_1c_ °C ± SEM) were comparable between species (excluding RCNT); BHNT (winter = 19.4 ± 2.7 °C, summer = 23.3 ± 3.14 °C), CCBF (19.2 ± 2.46 °C), and GBWW (22.3 ± 3.06 °C) (see Fig. 1). Similarly, estimates of thermal conductance (W °C ± SEM) were also comparable between species; BHNT (winter= 0.0127 ± 0.004 °C W, summer= 0.0156 ± 0.005 °C W), CCBF (0.0120 ± 0.004 °C W) and GBWW (0.0097 ± 0.002 °C W). No significant differences in either T_1c_ or C_min_ were detected between seasons for BHNT (T_1c_: *T* = 1.2141, df = 57, *P* = 0.229; C_min_: *T* = 0.6555, df = 57 *P* = 0.514), but BMR was significantly different between seasons (*T* = 2.3876, df = 57, *P* = 0.02). For full non-linear mixed model fits see Table S5.

In BHNT, summer BMR (0.43 ± 0.05 W) was 19.4% higher than winter BMR (0.36 ± 0.07 W). No seasonal differences in BMR were detected in CCBF (summer = 0.53 ± 0.04 W / winter = 0.52 ± 0.05 W) or GBWW (summer = 0.27 ± 0.04 W / winter = 0.31W), (Table 1, Fig. 1), although the winter sample for the latter species is from a single bird.

**Table.** Error! No text of specified style in document.. Seasonal comparisons of body mass (g), basal metabolic rate (BMR, whole animal (W) and mass-corrected (mW g^-1^)), and total blood haemoglobin content (g/dL) per species. Sample sizes (number of unique individuals) are presented in parentheses. Statistically significant differences between seasons are denoted with an asterisk. Note, samples sizes for BMR are a subset of the full dataset displayed in Fig. 1, where BMR is taken only from measurements greater than the lower critical temperature.

|  | Winter | Summer | W / S ratio |
| --- | --- | --- | --- |
| **Black-headed nightingale-thrush** *Catharus mexcanus* |  |  |  |
| Body Mass (g) | 30.40 ± 1.28 (20) | 31.04 ± 1.45 (64) | 0.98 |
| Whole-animal BMR (W) | 0.36 ± 0.07 (14) * | 0.43 ± 0.05 (15) * | 0.83 |
| Mass-specific BMR (mW g<sup>-1</sup>) | 12.02 ± 2.28 (14) * | 13.88 ± 1.95 (15) * | 0.87 |
| Total blood haemoglobin (g/dL) | 13.73 ± 1.00 (20) | 13.85 ± 1.06 (62) | 0.99 |
| **Ruddy-capped nightingale-thrush** *Catharus frantzii* |  |  |  |
| Body Mass (g) | - | 28.22 ± 0.95 (11) | - |
| Whole-animal BMR (W) | - | 0.47 ± 0.09 (11) | - |
| Mass-specific BMR (mW g<sup>-1</sup>) | - | 16.77 ± 3.50 (11) | - |
| Total blood haemoglobin (g/dL) | - | 15.53 ± 1.05 (11) | - |
| **Chestnut-capped brushfinch** *Arremon brunneinucha* |  |  |  |
| Body Mass (g) | 42.75 ± 2.17 (14) | 42.32 ± 1.55 (13) | 1.01 |
| Whole-animal BMR (W) | 0.52 ± 0.05 (8) | 0.53 ± 0.04 (5) | 0.98 |
| Mass-specific BMR (mW g<sup>-1</sup>) | 12.24 ± 1.46 (8) | 12.64 ± 1.05 (5) | 0.96 |
| Total blood haemoglobin (g/dL) | 13.55 ± 0.98 (14) | 13.8 ± 1.06 (13) | 0.99 |
| **Grey-breasted wood wren** *Henicorhina leucophrys* |  |  |  |
| Body Mass (g) | 15.35 ± 0.21 (2) * | 15.97 ± 0.60 (12) * | 0.96 |
| Whole-animal BMR (W) | 0.31 (1) | 0.27 ± 0.04 (4) | 1.14 |
| Mass-specific BMR (mW g<sup>-1</sup>) | 20.53 (1) | 17.03 ± 2.78 (4) | 1.2 |
| Total blood haemoglobin (g/dL) | 13.33 ± 0.94 (2) | 13.03 ± 0.72 (11) | 1.02 |
| **Common bush tanager** *Chlorospingus flavopectus* |  |  |  |
| Body Mass (g) | 15.8 ± 0.89 (38) * | 16.85 ± 0.45 (17) * | 0.93 |
| Total blood haemoglobin (g/dL) | 13.95 ± 1.02 (38) | 13.59 ± 1.05 (17) | 1.02 |

BMR in our species was generally greater than predicted (mean % difference ± SD) when compared to mass-scaling exponents for tropical birds. Winter BMR for BHNT (111.5 ± 3) was within the expected range, whereas summer BMR for BHNT (129.8 ± 3.5), summer RCNT (150.4 ± 2.9), and pooled values across season for CCBF (132.1 ± 3.7) and GBWW (125 ± 1) were greater than expected (Fig. 2).

**Fig. 2.**
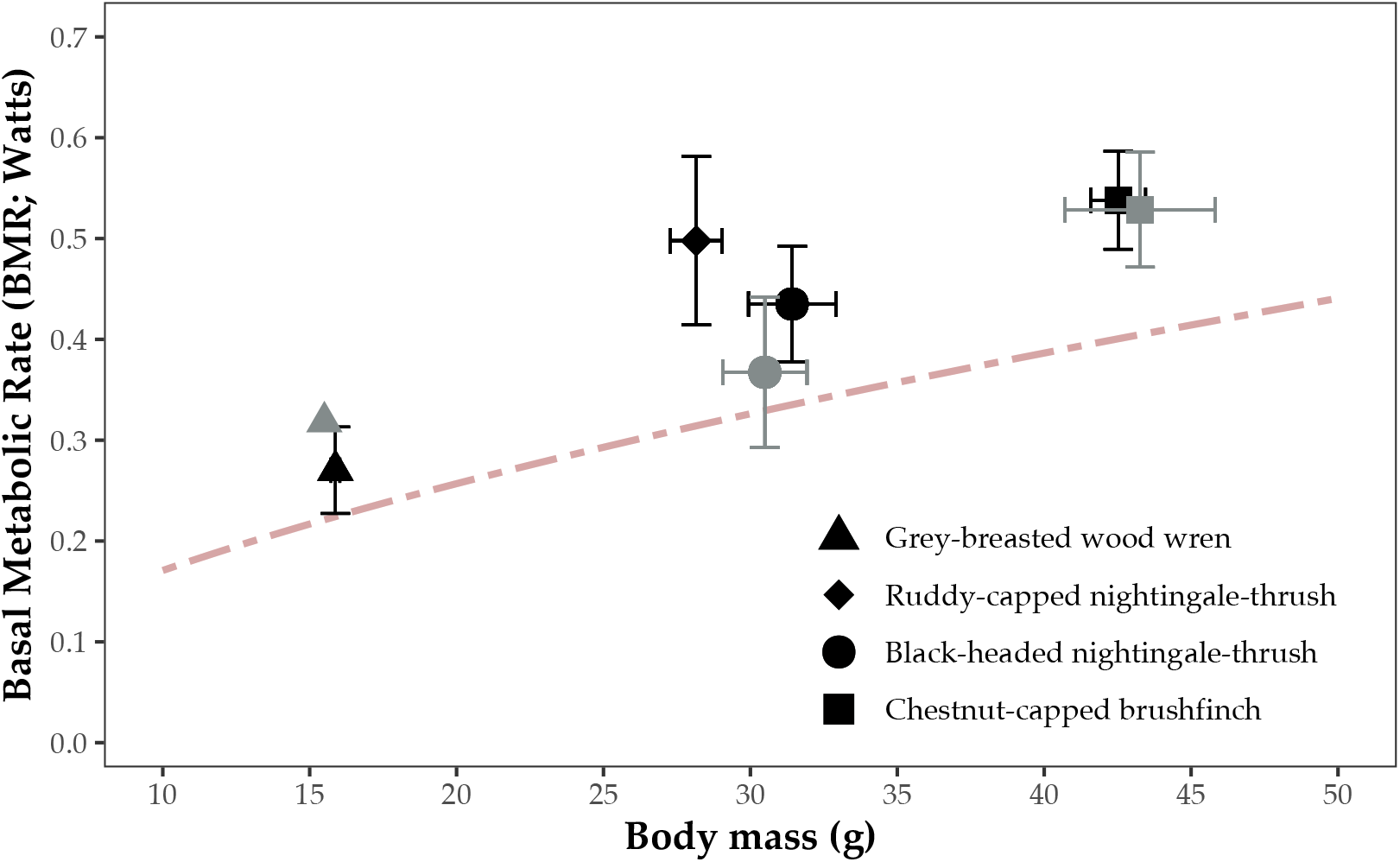
Mean (± SD) basal metabolic rate (BMR; W) and body mass (M_b_; g) per species between summer (black) and winter (grey). Although the only significant difference in metabolic rates between season was in black-headed nightingale-thrushes *Catharus mexicanus*, we plot all data separately for purposes of visual comparison. Expected allometric scaling relationship between basal metabolic rate and body mass for tropical birds (red dashed line) is taken from Londoño *et al*. (2015).

### Body mass

We detected seasonal changes in M_b_ in two of the four species for which we had data (Fig. 2, Table 1). No seasonal changes in M_b_ were evident in BHNT and CCBF (BHNT, χ^2^_1_ =3.06, *P* = 0.08; CCBF, χ^2^_1_ = 0.49, *P* = 0.48), whereas GBWW and COBT had higher M_b_ in the breeding season (GBWW, χ^2^_1_ = 222.8, *P* <0.01; COBT, χ^2^_1_ = 21.09, *P* <0.001). In addition, for BHNT we found no seasonal changes in M_b_ dependent on sex (χ^2^_1_ = 2.27, *P* = 0.13).

### Haemoglobin concentrations

In all study species, H_b_ concentrations varied little with changes in elevation, between season, and between sexes, with intercept only models performing best for each species. For BHNT, however, there was evidence for an effect of season dependent on sex, such that H_b_ is slightly higher in summer, over winter males (the breeding season × sex interaction term was significant in the second ranked model for this species: χ^2^_1_ = 4.923, *P* = 0.03; Table S6). For GBWW, elevation was also significant in the second ranked model (χ^2^_1_ = 5.074, *P* = 0.02; Table S9), suggesting that H_b_ decreases with increases in elevation (although the limited sample may explain this effect). Overall, however, H_b_ concentrations for each of the study species did not appear to be influenced by any of the hypothesised variables (for all species model fits see Tables S6-S10).

Blood H_b_ concentrations were comparable between all species, excepting RCNT (Fig. 3, Table 1). We found no intraspecific differences in winter H_b_ concentrations (all species comparisons *P* ≥ 0.55). Similarly, we also found no intraspecific differences in summer H_b_ concentrations (*P* ≥ 0.10) except for RCNT, which was distinctly higher than the other four species (*P* <0.001 for all RCNT – other species comparisons). For all post-hoc comparison tests see Table S11.

**Fig. 3.**
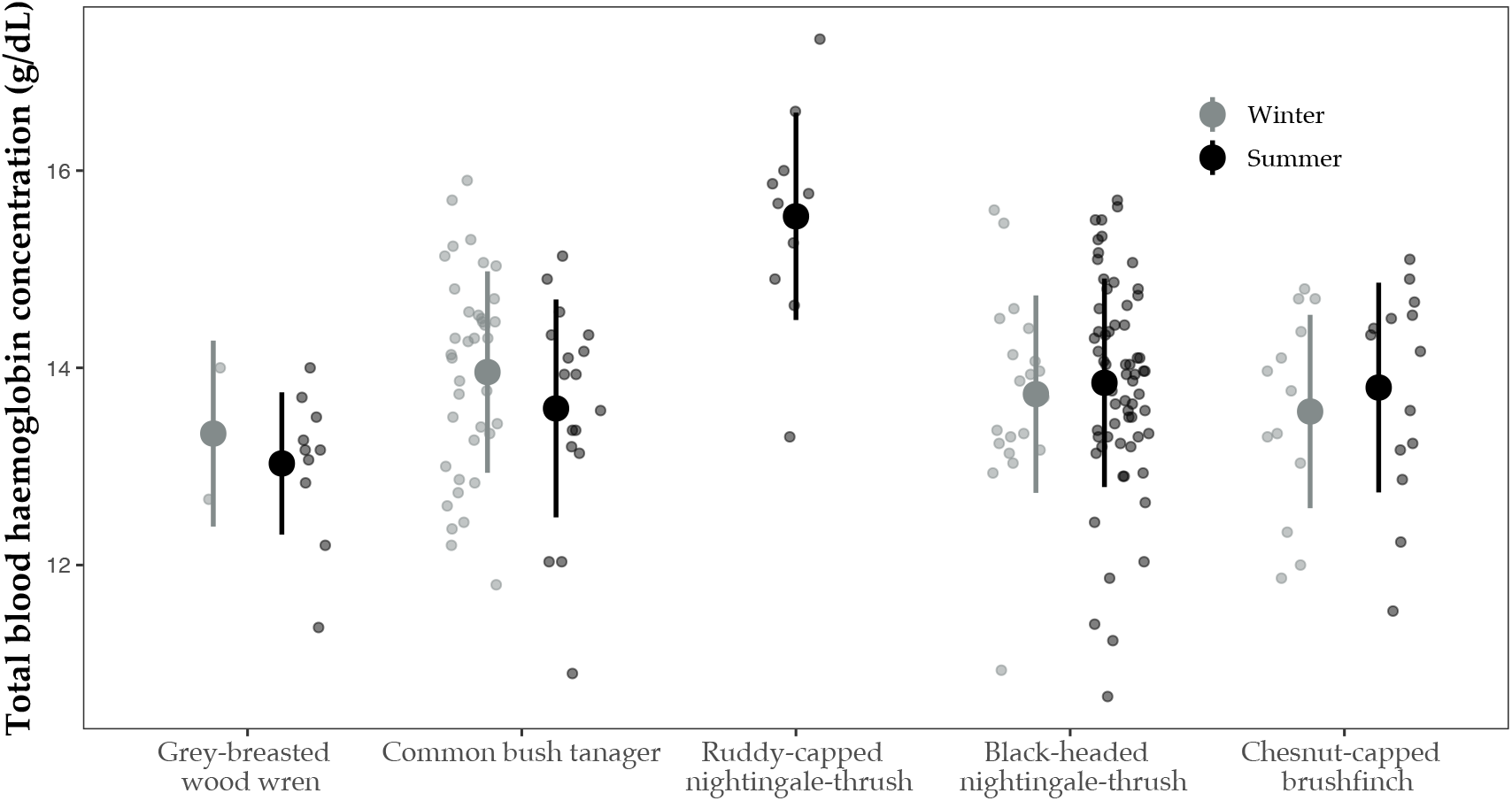
Haemoglobin concentrations (g/dL, mean ± SD) between species and between season. No winter data were available for ruddy-capped nightingale-thrushes *Catharus frantzii* (see Methods and Materials).

## Discussion

The aims of our study were twofold. Firstly, we wanted to determine whether our study species displayed uniform shifts in physiological traits across season, and secondly, to ascertain whether there was evidence for cold adaption in a higher elevation tropical species. Contrary to our first hypothesis, for the three species (BHNT, CCBF and GBWW) for which we had data between seasons, only BHNT regulated metabolic rates between season. This species up-regulated BMR in the warmer summer, whereas CCBF and GBWW showed no evidence for seasonal changes (although we recognise the sample size for winter GBWW is small and possibly inconclusive). We also found no evidence for seasonal shifts in H_b_ concentrations and variable seasonal changes in M_b_. In support of our second hypothesis, however, RCNT was a clear outlier in all measures, with evidence of cold tolerance traits (no discernible metabolic response to temperature), higher summer BMR relative to M_b_, and higher H_b_ concentration than other species.

### Physiological shifts with season

The variability in metabolic rate between seasons observed across species in our study is consistent with comparable studies on lowland tropical birds. For example, the interspecific variation in BMR in our study (W/S ratio range 0.83-1.14; Table 1) is within the range of that measured in lowland tropical species in Panama (W/S ratio range 0.71-1.33; Pollock *et al.* (2019)). Because our study was at higher elevation, and thus a cooler T_e_, we hypothesised that seasonal changes in metabolic rates may reflect these cooler T_e_ and minor changes in season. Instead, our results appear unrelated to T_e_ because no shifts in metabolic rates were consistent with heat conservancy (i.e. increases in BMR and thermoregulatory traits in winter) as is common among temperate species (Swanson & Garland 2009, Swanson 2010, Smit & McKechnie 2010, McKechnie *et al.* 2015). This apparent lack of relationship with T_e_ was also reflected in M_b_ between season, which were consistent, or increased in summer (GBWW and COBT), also opposite of seasonal changes in M_b_ typical among temperate species. In a similar study, Pollock *et al*. (2019) did not detect any significant differences in M_b_ changes between temperate and tropical birds, although these authors also found substantial variation among tropical species, similar to our results.

Our study is only the third we are aware of to compare metabolic rates between seasons in tropical species. Like other studies, however, we also show that variation in metabolic rate in tropical birds is apparently unrelated to T_e_ and the direction of change is highly variable across species (Wells & Schaeffer 2012, Pollock *et al.* 2019). Although there are few studies from the tropics to which we can make direct comparisons, this variation is also consistent with seasonal changes in metabolic rates in subtropical birds. Substantial variation in BMR has also been documented in subtropical species, ranging from considerable winter reductions, to considerable winter increases (W/S ratio range 0.66-1.63; McKechnie *et al.* 2015), a magnitude of shifts comparable to temperate species but with less predictable seasonal directionality (Noakes *et al.* 2017).

Because seasonal changes in metabolic rates appears unrelated to T_e_ in our, and other, studies in the tropics, what drives these seasonal regulations (or lack thereof) remains unclear. One emerging explanation for this pattern is that ‘metabolic niches’ are greater at tropical latitudes (Anderson & Jetz 2005), where intrinsic relationships between species characteristics and metabolic rate reflect the seasonal variation in metabolic rates (McKechnie *et al.* 2015). For example, specific behavioural changes with season unique to a given species may manifest in changing maintenance costs of metabolically active tissues (Swanson 2010). In BHNT, BMR was up-regulated in summer (19.4% greater than winter), the opposite of patterns typically displayed in temperate species (e.g. Smit & McKechnie 2010). Thus, behavioural differences related to reproductive activity between season in our study species may covary with BMR because of the energetic costs associated with them (e.g. increased activity rates in males). That physiological control mechanisms may shape the diversity of life history traits in tropical birds has previously been suggested (Ricklefs & Wikelski 2002, Williams *et al.* 2010). However, we are aware of only one study that has assessed the relationship between metabolic rate and behavioural energy usage in a tropical bird (Steiger *et al.* 2009), and none that have assessed seasonal differences in metabolic intensity with lifehistory traits in tropical species.

A shortfall in our interpretation of the processes underpinning these seasonal changes, however, is that we did not concurrently measure body temperatures. Facultative hyperthermia, where short-term reductions in body temperature reduce the energetic demands of metabolic heat production are thought to be common across birds (McKechnie & Lovegrove 2002) and have been found in various tropical species, often concurrently with reductions in metabolic rate (Bartholomew et al. 1983, Merola-Zwartjes 1998, Merola-Zwartjes & Ligon 2000, Steiger et al. 2009, Burnett et al. 2019). As such, we cannot eliminate seasonal changes in body temperature as an alternative explanation for the interspecific differences we observe. Irrespective of the mechanism, however, our results still display species-specific changes in metabolic rate between seasons.

That we found no clear changes in H_b_ concentrations in our study species with season is intriguing, particularly when coupled with the variation observed in BMR. Firstly, thermogenic demands may result in elevated H_b_ concentrations (Swanson 1990, Powell *et al.* 2013), so our result that H_b_ concentrations appeared unchanged between seasons is consistent with the lack of other apparent physiological changes related to increased cold tolerance. Secondly, because H_b_ concentrations reflect oxidative stress (Minias 2015), the lack of seasonal change in H_b_ concentrations in the species in our study are probably indicative of limited physiological stress between seasons. This may be particularly the case in BHNT, where seasonal changes in BMR did not appear to covary with H_b_ concentrations, suggesting that increased physiological stress during reproduction is not a sufficient explanation for seasonal changes in BMR observed in this species, at least in males. We remain cautious in this conclusion, however, because there was some signal of a difference in male H_b_ concentrations between season, although there was much overlap and quantities were similar (H_b_ g/dL content in winter (n = 15)/summer (n = 54) male BHNT; 13.5 ± 0.98 / 13.9 ± 1.05).

We are aware of no other studies on H_b_ concentrations in tropical forest birds from which to directly compare, but the general lack of seasonal changes across our study species may be reflective of ‘slow paced’ life-histories of tropical birds, where longevity is facilitated by greater investment in self-maintenance (Ricklefs & Wikelski 2002, Wiersma *et al.* 2007). Although, only on temperate species, evidence in support of a coevolution with slower paced life-history and decreased levels of oxidative stress has been displayed (Vágási *et al.* 2018). Reproductive periods in female birds may be particularly energetically demanding (Williams *et al.* 2004) but because of the difficulties in determining sex in many of the species in our study in the winter, we were not able to fully interrogate this hypothesis. Future studies could clarify this by comparing magnitudes of change between tropical and temperate species with the addition of new field data (which for H_b_ concentrations are simple and cost-effective to obtain).

### Implications for elevational adaption

Among our study species, RCNT was distinct. We found no low temperature limit at which this species began thermoregulation, a higher BMR than predicted by mass-scaling exponents among the species in our study (over greater 50% greater than that predicted for tropical birds), and higher H_b_ concentration than other study species: all traits consistent with cold tolerance (Swanson 2010, Pollock *et al.* 2019). Few studies on the physiology of tropical montane birds exist for which we can directly compare, although similar (albeit isolated) examples of cold tolerant birds in the Peruvian Andes (Londoño *et al.* 2017) and high elevation tropical bats and mice have been found (Soriano *et al.* 2002, Pasch *et al.* 2013).

The physiological differences apparent in RCNT are particularly intriguing when considered against the backdrop of elevational range restrictions characteristic of tropical montane species (e.g. McCain 2009). Realistically, it is unlikely that the traits we observe are a product of differences in present day T_e_ across elevation in our site; T_e_ did not strikingly differ across elevation, and that BHNT, a parapatric congener (see Jones *et al.* 2019) displays divergent physiological traits. The two species of nightingale-thrush are not sister species (Voelker *et al.* 2013) and one possibility is that the divergence in RCNT is a product of conserved traits from historic isolation (Wiens *et al.* 2010). That the species are parapatric and compete at their elevational range limits (Jones *et al.* 2019) has likely developed as a result of range convergence (Freeman 2015). This trait divergence (and competitive interactions) resembles that of morphological phenotypes between two parapatric wood wrens (*Henicorhina* sp.) in Colombia, the elevational ranges of which have converged through secondary contact (Caro *et al.* 2013). Further supporting this hypothesis, elevational range has been suggested as an important driver of the evolution in variation of blood oxygen carrying capacity in birds (Minias 2020).

A general lack of evidence for intraspecific variation in H_b_ concentrations or metabolic rate with increases in elevation is consistent with a lack of apparent elevational specialism in our study species (excepting RCNT). However, because we only tested for an intraspecific effect across a small elevational range we remain cautious in this conclusion as this may not be conclusive evidence that one does not exist. Intraspecific differences ascribed to elevational adaption have been displayed in Tyrannid flycatchers (*Anairetes* sp.) across a larger elevational gradient (~1200m) in the Peruvian Andes (Dubay & Witt 2014) and for resident species in the subtropical Himalayas (Barve *et al.* 2016). Nonetheless, a lack of intraspecific signal in physiological traits across elevation is still consistent with an apparent lack of elevational specialism, although more comprehensive tests across a broader elevational range at our study site would clarify this, particularly with the inclusion of lower elevation species.

The species in our study had generally higher BMR than predicted by M_b_ for tropical birds, with only winter BMR of BHNT falling within the expected values. To some extent this contrasts with Londoño *et al.* (2015) who, across a large range of species, displayed that higher elevation tropical species had comparable BMR to lowland tropical residents. The specific reasons our BMR measures were slightly higher than predicted is not clear and did not appear to be a result of intraspecific variation across elevation (e.g. Lindsay *et al.* 2012). One possibility is that species that do not undergo seasonal shifts in BMR maintain overall higher BMR throughout the year, although because little is known of the specific drivers of seasonal shifts in BMR in tropical species, rigorous testing is required to determine this. Nonetheless, despite generally higher BMR than predicted by allometric scaling, our values are still within the range previously measured for tropical birds (Fig. S2).

In conclusion, our results support two emergent patterns in our understanding of the physiological diversity of tropical birds. Firstly, that seasonal changes in metabolic rates appear to be flexible and possibly species specific, more broadly reflecting the growing appreciation for flexible phenotypic diversity in metabolic rates in birds (Piersma & Drent 2003, McKechnie *et al.* 2006, McKechnie 2007). Secondly, that the physiology of tropical birds appears unrelated to T_e_ as conventional hypotheses have suggested (see Chown *et al.* 2004, Londoño *et al.* 2015). However, we did find evidence of distinct interspecific differences in our study, the generalities of which require more comprehensive examination. Despite growing interest, substantial knowledge gaps remain in our understanding of the ecophysiology of tropical birds. This is particularly so for tropical montane species, that may be characterised by distinct interplays between physiological and life-history characteristics (Goymann *et al.* 2004, Scholer *et al.* 2019).

## Acknowledgements

This study would not have been possible without the logistical support of numerous staff and volunteers of Operation Wallacea in Cusuco National Park, Honduras. All work was undertaken under permits issued by Instituto de Conservacion Forestal in Honduras, and with ethical approval by Royal Holloway University of London ethics committee. We also thank Craig White for technical support. This work was supported by the Natural Environment Research Council [grant number NE/L002485/1].

