## Supplemental materials for "Comparative physiology of five tropical montane songbirds reveals differential seasonal acclimatisation and cold adaption"

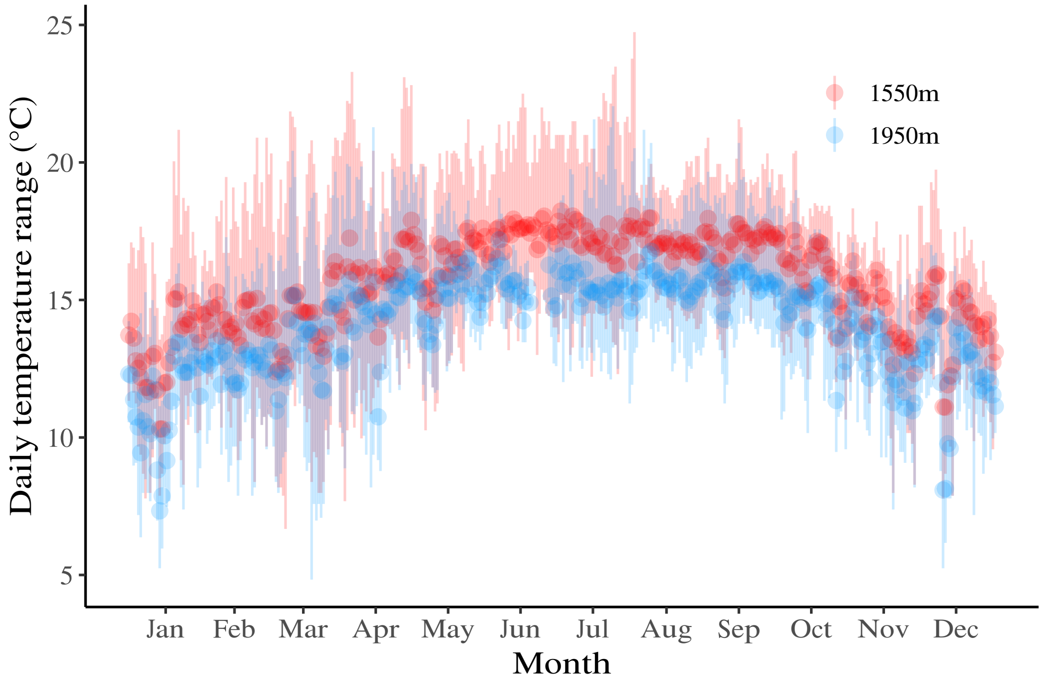

**Fig. S1.** Daily mean (points) and range of minimum and maximum (lines) environmental temperature (°C) variation across the year at the two research camps (1550m and 1950m, respectively) in Cusuco National Park, Honduras.

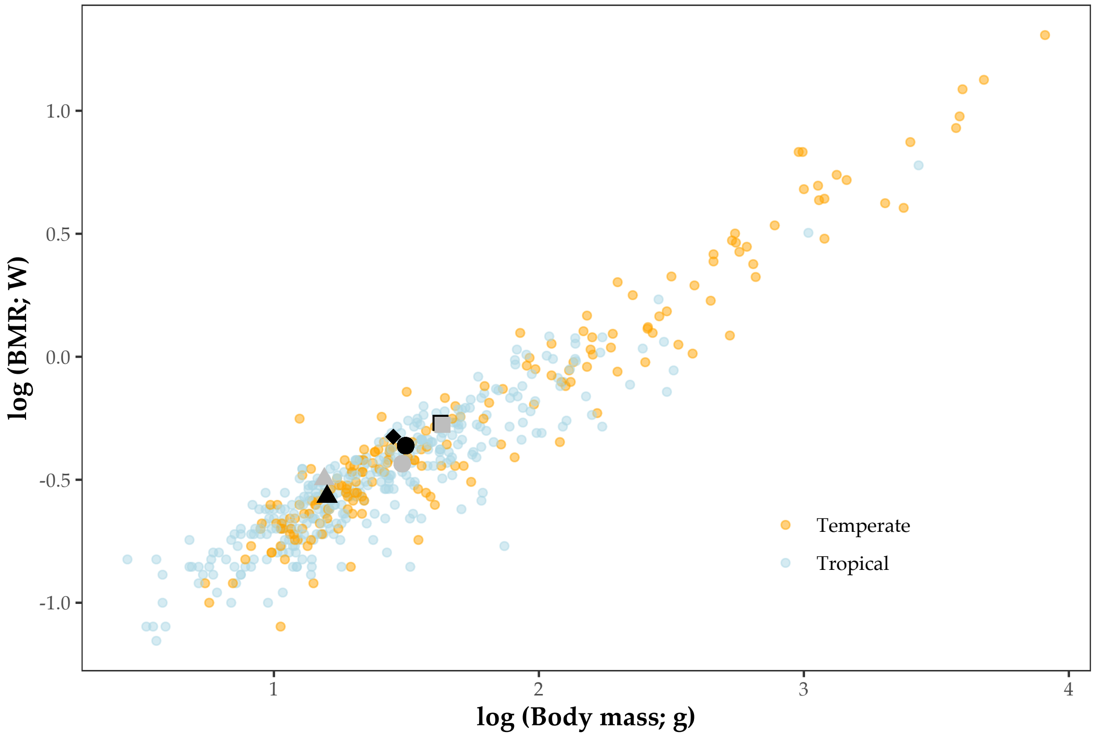

**Fig. S2** Basal metabolic rate (BMR; Watts) as a function of body mass (g) (both log-transformed) for tropical and temperate birds using data from Londoño *et al*. (2015a, b). Species in this study (symbols as per Fig. 2 main text) are split for summer (black) and winter (grey) for purposes of visual comparison to global BMR data. One extreme outlier was removed from this dataset (common ostrich *Struthio camelus*) for better visual comparison.

**Table S**Error! No text of specified style in document.**.** Linear mixed-effects model selections for metabolic rate (W) as a function of treatment temperature, breeding season, body mass (g) and elevation (m) for black-headed nightingale-thrushes *Catharus mexicanus* (A) and repeated model selections on mass-corrected values (B). Model fits are presented as a full model (inclusive of all parameters), null models (intercept only) and best model fits as ranked by AICc scores. The best fitting model is denoted with Δ0, with all models within Δ6 AICc also presented. Parsimonious models of significant variables are also presented from model fits where best fitting models had multiple parameters. Statistically significant effects are assessed by Wald $\chi^2$ tests. The 'BreedingY' parameter represents the summer breeding season.

**A**

| | AICc | df | Variable | Estimate | SE | *t*-value | $x^2$ | df | *p*-value |
|---|---|---|---|---|---|---|---|---|---|
| Full Model | -189.4 | 8 | Intercept | 0.4408 | 0.0138 | 31.88 | | | |
| | | | Temperature | -0.0419 | 0.0100 | -4.16 | 66.444 | 1 | **<0.001** |
| | | | BreedingY | 0.0697 | 0.0172 | 4.05 | 16.999 | 1 | **<0.001** |
| | | | Mass | 0.0235 | 0.0085 | 2.76 | 7.619 | 1 | **<0.01** |
| | | | Elevation | 0.0021 | 0.0082 | 0.25 | 0.064 | 1 | 0.798 |
| | | | Temperature* Breeding | -0.0333 | 0.0143 | -2.32 | 5.396 | 1 | **<0.05** |
| Null Model | -165.6 | 3 | Intercept | 0.4832 | 0.0122 | 39.51 | | | |
| Best model | -206.5 (Δ0) | 5 | Intercept | 0.4324 | 0.0146 | 29.58 | | | |
| | | | Temperature | -0.0564 | 0.0073 | -7.66 | 58.708 | 1 | **<0.001** |
| | | | BreedingY | 0.0828 | 0.0177 | 4.68 | 21.872 | 1 | **<0.001** |
| | -203.1 (Δ3.39) | 6 | Intercept | 0.4412 | 0.0146 | 30.19 | | | |
| | | | Temperature | -0.0587 | 0.0072 | -8.13 | 66.065 | 1 | **<0.001** |
| | | | BreedingY | 0.0691 | 0.0180 | 3.84 | 14.707 | 1 | **<0.001** |
| | | | Mass | 0.0230 | 0.0089 | 2.57 | 6.618 | 1 | **<0.05** |
| | -201.9 (Δ4.59) | 6 | Intercept | 0.4324 | 0.0139 | 31.02 | | | |
| | | | Temperature | -0.0408 | 0.0103 | -3.95 | 59.448 | 1 | **<0.001** |
| | | | BreedingY | 0.0831 | 0.0170 | 4.87 | 23.978 | 1 | **<0.001** |
| | | | Temperature* BreedingY | -0.0309 | 0.0146 | -2.12 | 4.479 | 1 | **<0.05** |
| *Parsimonious models* | | | | | | | | | |
| | 196.2 (Δ10.24) | 4 | Intercept | -0.4857 | 0.0113 | 42.87 | | | |
| | | | Temperature | -0.0567 | 0.0078 | -7.24 | 52.352 | 1 | **<0.001** |
| | -171.2 (Δ35.31) | 4 | Intercept | 0.4317 | 0.0168 | 25.66 | | | |
| | | | BreedingY | 0.0816 | 0.0208 | 3.91 | 15.268 | 1 | **<0.001** |

**B** (Mass-corrected values)

| | | | | | | |
|---|---|---|---|---|---|---|
| Full | -864 | 7 | Intercept | 0.0508 | 0.0015 | 32.98 |

| | | | | | | | | | |
|---|---|---|---|---|---|---|---|---|---|
| Model | | | Temperature | -0.0049 | 0.0011 | -4.24 | 66.698 | 1 | **<0.001** |
| | | | BreedingY | 0.0083 | 0.0018 | 4.39 | 19.510 | 1 | **<0.001** |
| | | | Elevation | 0.0003 | 0.0009 | 0.34 | 0.112 | 1 | 0.737 |
| | | | Temperature* Breeding | -0.0035 | 0.0016 | -2.17 | 4.691 | 1 | **<0.05** |
| Null Model | -863.7 | 3 | Intercept | 0.0559 | 0.0013 | 40.99 | | | |
| Best model | -893 (Δ0) | 4 | Intercept | 0.0157 | 0.0003 | 45.76 | | | |
| | | | Temperature | -0.0018 | 0.0002 | -7.81 | 60.974 | 1 | **<0.001** |
| | -892.5 (Δ0.47) | 5 | Intercept | 0.0142 | 0.0005 | 31.38 | | | |
| | | | Temperature | -0.0018 | 0.0002 | -8.16 | 66.580 | 1 | **<0.001** |
| | | | BreedingY | 0.0022 | 0.0005 | 4.15 | 17.284 | 1 | **<0.001** |

**Table S2.** Linear mixed-effects model selections for metabolic rate (W) as a function of treatment temperature, body mass and elevation for ruddy-capped nightingale-thrushes *Catharus frantzii*. Model fits are presented as a full model (inclusive of all parameters), null models (intercept only) and best model fits as ranked by AICc scores. The best fitting model is denoted with Δ0, with all models within Δ6 AICc also presented. Statistically significant effects are assessed by Wald $\chi^2$ tests.

| | AICc | df | Variable | Estimate | SE | *t*-value | $x^2$ | df | *p*-value |
|---|---|---|---|---|---|---|---|---|---|
| Full Model | -21.5 | 6 | Intercept | 0.4979 | 0.0150 | 33.09 | | | |
| | | | Temperature | -0.0256 | 0.0171 | -1.49 | 2.227 | 1 | 0.135 |
| | | | Mass | 0.0061 | 0.0169 | 0.36 | 0.130 | 1 | 0.717 |
| | | | Elevation | -0.0366 | 0.0163 | -2.25 | 5.044 | 1 | **<0.05** |
| Null Model | -42.9 (Δ0) | 3 | Intercept | 0.4979 | 0.0163 | 30.42 | | | |
| | -37.5 (Δ5.39) | 4 | Intercept | 0.4979 | 0.0154 | 32.22 | | | |
| | | | Elevation | -0.0317 | 0.0157 | -2.01 | 4.056 | 1 | **<0.05** |

**Table S3.** Linear mixed-effects model selections for metabolic rate (W) as a function of treatment temperature, breeding season, body mass and elevation for chestnut-capped brushfinches *Arremon brunneinucha* (A) and repeated model selections on mass-corrected values (B). Model fits are presented as a full model (inclusive of all parameters), null models (intercept only) and best model fits as ranked by AICc scores. The best fitting model is denoted with Δ0, with all models within Δ6 AICc also presented. Statistically significant effects are assessed by Wald $\chi^2$ tests. The 'BreedingY' parameter represents the summer breeding season.

**A**

| | AICc | df | Variable | Estimate | SE | *t*-value | $x^2$ | df | *p*-value |
|---|---|---|---|---|---|---|---|---|---|
| Full Model | -47.7 | 8 | Intercept | 0.5507 | 0.0158 | 34.68 | | | |
| | | | Temperature | -0.0506 | 0.0114 | -4.41 | 28.859 | 1 | **<0.001** |
| | | | BreedingY | 0.0313 | 0.0271 | 1.15 | 1.543 | 1 | 0.214 |
| | | | Mass | 0.0215 | 0.0089 | 2.40 | 5.781 | 1 | **<0.05** |
| | | | Elevation | -0.0107 | 0.0146 | -0.73 | 0.535 | 1 | 0.464 |
| | | | Temperature* Breeding | 0.0029 | 0.0187 | 0.16 | 0.024 | 1 | 0.876 |
| Null Model | -72.2 | 3 | Intercept | 0.5656 | 0.0116 | 48.75 | | | |
| Best model | -81.4 (Δ0) | 4 | Intercept | 0.5688 | 0.0099 | 57.05 | | | |
| | | | Temperature | -0.0466 | 0.0086 | -5.40 | 29.19 | 1 | **<0.001** |
| | -75.1 (Δ6.29) | 5 | Intercept | 0.5663 | 0.0086 | 65.40 | | | |
| | | | Temperature | -0.0477 | 0.0084 | -5.66 | 32.034 | 1 | **<0.001** |
| | | | Mass | 0.019 | 0.0088 | 2.17 | 4.687 | 1 | **<0.05** |

**B** (mass-corrected values)

| | AICc | df | Variable | Estimate | SE | *t*-value | $x^2$ | df | *p*-value |
|---|---|---|---|---|---|---|---|---|---|
| Full Model | -268.2 | 7 | Intercept | 0.0129 | 0.0003 | 35.62 | | | |
| | | | Temperature | -0.0012 | 0.0002 | -4.52 | 30.909 | 1 | **<0.001** |
| | | | BreedingY | 0.0007 | 0.0006 | 1.24 | 1.786 | 1 | 0.181 |
| | | | Elevation | -0.0002 | 0.0003 | -0.81 | 0.660 | 1 | 0.416 |
| | | | Temperature* Breeding | $5.939 \times 10^{-5}$ | 0.0004 | 0.13 | 0.018 | 1 | 0.892 |
| Null Model | -311.8 (Δ5.4) | 3 | Intercept | 0.0132 | 0.0002 | 48.46 | | | |
| Best model | -317.2 (Δ0) | 4 | Intercept | 0.0133 | 0.0001 | 67.13 | | | |
| | | | Temperature | -0.0011 | 0.0001 | -5.79 | 33.548 | 1 | **<0.001** |

**Table S4.** Linear mixed-effects model selections for metabolic rate (W) as a function of treatment temperature, breeding season, body mass and elevation for grey-breasted wood wrens *Henicorhina leucophrys*. Model fits are presented as a full model (inclusive of all parameters), null models (intercept only) and best model fits as ranked by AICc scores. The best fitting model is denoted with Δ0 (all other model iterations where greater than Δ6 AICc). Statistically significant effects are assessed by Wald $\chi^2$ tests. The 'BreedingY' parameter represents the summer breeding season.

| | AICc | df | Variable | Estimate | SE | *t*-value | $x^2$ | df | *p*-value |
|---|---|---|---|---|---|---|---|---|---|
| Full Model | -9.2 | 8 | Intercept | 0.2386 | 0.0608 | 3.92 | | | |
| | | | Temperature | 0.0678 | 0.0616 | 1.10 | 16.739 | 1 | **<0.001** |
| | | | BreedingY | 0.0979 | 0.0628 | 1.55 | 0.114 | 1 | 0.7357 |
| | | | Mass | 0.0108 | 0.0117 | 0.92 | 0.856 | 1 | 0.3538 |
| | | | Elevation | -0.0036 | 0.0122 | -0.30 | 0.091 | 1 | 0.763 |
| | | | Temperature* Breeding | -0.1169 | 0.0626 | -1.86 | 3.480 | 1 | 0.062 |
| Null Model | -40.4 (Δ3.93) | 3 | Intercept | 0.3354 | 0.0148 | 22.62 | | | |
| Best model | -44.3 (Δ0) | 4 | Intercept | 0.3354 | 0.0106 | 31.48 | | | |
| | | | Temperature | -0.0484 | 0.0109 | -4.44 | 19.720 | 1 | **<0.001** |

**Table S5.** Non-linear mixed models per species on metabolic rate (W) as a function of temperature estimating lower critical temperatures ($T_{lc}$), minimum thermal conductance ($C_{min}$) and basal metabolic rate (BMR) above inflection temperature (see Fig. 1 main text) for all species except ruddy-capped nightingale-thrushes *Catharus frantzii*. A multi-level non-linear mixed model with the two-way factor of 'Season' (summer or winter) was fitted for black-headed nightingale-thrushes *Catharus mexicanus* because significant differences in metabolic rates were detected between seasons (see Table S1).

| | Variable | Value | SE | df | t-value | *p* value |
|---|---|---|---|---|---|---|
| **Black-headed nightingale-thrush** *Catharus mexicanus* | | | | | | |
| *Winter* *n* = 15 | BMR (W) | 0.391 | 0.015 | 57 | 25.291 | **<0.001** |
| | $C_{min}$ (W °C) | 0.013 | 0.004 | 57 | 3.306 | **0.002** |
| | $T_{lc}$ (W °C) | 19.479 | 2.772 | 57 | 7.026 | **<0.001** |
| *Summer* *n* = 28 | BMR (W) | 0.441 | 0.022 | 57 | 2.388 | **0.02** |
| | $C_{min}$ (W °C) | 0.016 | 0.005 | 57 | 0.656 | 0.514 |
| | $T_{lc}$ (°C) | 23.303 | 3.149 | 57 | 1.214 | 0.229 |
| **Chestnut-capped brushfinch** *Arremon brunneinucha* | | | | | | |
| *n* = 14 | BMR (W) | 0.534 | 0.012 | 17 | 43.399 | **<0.001** |
| | $C_{min}$ (W °C) | 0.012 | 0.004 | 17 | 3.371 | **0.004** |
| | $T_{lc}$ (°C) | 19.238 | 2.469 | 17 | 7.791 | **<0.001** |
| **Grey-breasted wood wren** *Henicorhina leucophrys* | | | | | | |
| *n* = 8 | BMR (W) | 0.291 | 0.016 | 11 | 18.023 | **<0.001** |
| | $C_{min}$ (W °C) | 0.010 | 0.003 | 11 | 3.310 | **0.007** |
| | $T_{lc}$ (°C) | 22.308 | 3.069 | 11 | 7.268 | **<0.001** |

**Table S6** Linear mixed effect model selections for total blood haemoglobin content (g/dL) as a function of sex, season and elevation for black-headed nightingale-thrushes *Catharus mexicanus* as ranked by AICc scores. The best fitting model is denoted with Δ0, with all models within Δ6 AICc also presented. Statistically significant effects are assessed by Wald $\chi^2$ tests. The 'BreedingY' parameter represents the summer breeding season, and 'SexM' as male birds.

| | AICc | df | Variable | Estimate | SE | *t*-value | $x^2$ | df | *p*-value |
|---|---|---|---|---|---|---|---|---|---|
| Full Model | 256.3 | 7 | Intercept | 14.489 | 0.461 | 31.43 | | | |
| | | | Elevation | 0.038 | 0.115 | 0.33 | 0.109 | 1 | 0.741 |
| | | | SexM | -0.982 | 0.532 | -1.85 | 0.005 | 1 | 0.942 |
| | | | BreedingY | -1.019 | 0.571 | -1.78 | 0.151 | 1 | 0.697 |
| | | | SexM* BreedingY | 1.426 | 0.644 | 2.22 | 4.925 | 1 | **<0.05** |
| Null Model | 249.4 (Δ0) | 3 | Intercept | 13.832 | 0.118 | 117.70 | | | |
| | 251.6 (Δ2.17) | 6 | Intercept | 14.485 | 0.458 | 31.61 | | | |
| | | | SexM | -0.981 | 0.529 | -1.86 | 0.007 | 1 | 0.929 |
| | | | BreedingY | -1.005 | 0.567 | -1.76 | 0.167 | 1 | 0.682 |
| | | | SexM* BreedingY | 1.418 | 0.640 | 2.22 | 4.923 | 1 | **<0.05** |
| | 252.1 (Δ2.69) | 4 | Intercept | 13.846 | 0.286 | 48.47 | | | |
| | | | SexM | -0.016 | 0.314 | -0.05 | 0.003 | 1 | 0.959 |
| | 252.3 (Δ2.88) | 4 | Intercept | 13.753 | 0.233 | 59.00 | | | |
| | | | BreedingY | 0.103 | 0.265 | 0.39 | 0.151 | 1 | 0.698 |
| | 254.0 (Δ4.62) | 4 | Intercept | 13.831 | 0.118 | 117.02 | | | |
| | | | Elevation | 0.029 | 0.116 | 0.25 | 0.064 | 1 | 0.801 |
| | 255.0 (Δ5.6) | 5 | Intercept | 13.777 | 0.335 | 41.17 | | | |
| | | | SexM | -0.029 | 0.317 | -0.09 | 0.009 | 1 | 0.926 |
| | | | BreedingY | 0.105 | 0.269 | 0.39 | 0.152 | 1 | 0.697 |

**Table S7** Linear mixed effect model selections for total blood haemoglobin content (g/dL) as a function of sex and elevation for ruddy-capped nightingale-thrushes *Catharus frantzii* as ranked by AICc scores. The best fitting model is denoted with Δ0, with all models within Δ6 AICc also presented. Statistically significant effects are assessed by Wald $\chi^2$ tests.

| | AICc | df | Variable | Estimate | SE | *t*-value | $x^2$ | df | *p*-value |
|---|---|---|---|---|---|---|---|---|---|
| Full Model | 43.6 | 4 | Intercept | 15.422 | 0.367 | 42.05 | | | |
| | | | Elevation | -0.124 | 0.361 | -0.34 | 0.117 | 1 | 0.732 |
| Null Model | 38.2 (Δ0) | 3 | Intercept | 15.439 | 0.343 | 45.02 | | | |

**Table S8** Linear mixed effect model selections for total blood haemoglobin content (g/dL) as a function of sex, season and elevation for chestnut-capped brushfinch *Arremon brunneinucha* as ranked by AICc scores. The best fitting model is denoted with Δ0, with all models within Δ6 AICc also presented. Statistically significant effects are assessed by Wald $\chi^2$ tests. The 'BreedingY' parameter represents the summer breeding season, and 'SexM' as male birds and 'SexU' as birds of undetermined sex.

| | AICc | df | Variable | Estimate | SE | *t*-value | $x^2$ | df | *p*-value |
|---|---|---|---|---|---|---|---|---|---|
| Full Model | 99.7 | 9 | Intercept | 13.959 | 0.526 | 26.55 | | | |
| | | | Elevation | -0.163 | 0.260 | -0.63 | 0.396 | 1 | 0.529 |
| | | | SexM | -0.652 | 0.652 | -0.99 | 0.071 | 2 | 0.965 |
| | | | SexU | -0.762 | 0.794 | -0.96 | | | |
| | | | BreedingY | -0.941 | 0.806 | -1.17 | 0.246 | 1 | 0.620 |
| | | | SexM* BreedingY | 1.667 | 0.949 | 1.76 | 4.279 | 2 | 0.118 |
| | | | SexU* BreedingY | 2.763 | 1.542 | 1.79 | | | |
| Null Model | 87.5 (Δ0) | 3 | Intercept | 13.679 | 0.191 | 71.59 | | | |
| | 90 (Δ2.41) | 4 | Intercept | 13.557 | 0.273 | 49.61 | | | |
| | | | Breeding | 0.243 | 0.387 | 0.63 | 0.395 | 1 | 0.530 |
| | 91.6 (Δ4.06) | 4 | Intercept | 13.679 | 0.194 | 70.38 | | | |
| | | | Elevation | 0.061 | 0.198 | 0.31 | 0.097 | 1 | 0.755 |
| | 92.2 (Δ4.64) | 5 | Intercept | 13.865 | 0.295 | 46.97 | | | |
| | | | SexM | -0.063 | 0.323 | -0.19 | 0.121 | 2 | 0.9412 |
| | | | SexU | 0.070 | 0.657 | 0.11 | | | |

**Table S9** Linear mixed effect model selections for total blood haemoglobin content (g/dL) as a function of season and elevation for grey-breasted wood wren *Henicorhina leucophrys* as ranked by AICc scores. The best fitting model is denoted with Δ0, with all models within Δ6 AICc also presented. Statistically significant effects are assessed by Wald $\chi^2$ tests. The 'BreedingY' parameter represents the summer breeding season.

| | AICc | df | Variable | Estimate | SE | *t*-value | $x^2$ | df | *p*-value |
|---|---|---|---|---|---|---|---|---|---|
| Full Model | 43.7 | 5 | Intercept | 12.891 | 0.504 | 25.57 | | | |
| | | | Elevation | -0.442 | 0.209 | -2.12 | 4.472 | 1 | **<0.05** |
| | | | BreedingY | 0.220 | 0.557 | 0.39 | 0.156 | 1 | 0.693 |
| Null Model | 37.4 (Δ0) | 3 | Intercept | 13.077 | 0.200 | 65.36 | | | |
| | 38.9 (Δ1.42) | 4 | Intercept | 13.077 | 0.173 | 75.64 | | | |
| | | | Elevation | -0.405 | 0.180 | -2.25 | 5.074 | 1 | **<0.05** |
| | 40.8 (Δ3.35) | 4 | Intercept | 13.333 | 0.526 | 25.34 | | | |
| | | | BreedingY | -0.303 | 0.572 | -0.53 | 0.281 | 1 | 0.596 |

**Table S10** Linear mixed effect model selections for total blood haemoglobin content (g/dL) as a function of sex, season and elevation for common bush tanager *Chlorospingus flavopectus* as ranked by AICc scores. The best fitting model is denoted with Δ0, with all models within Δ6 AICc also presented. Statistically significant effects are assessed by Wald $\chi^2$ tests. The 'BreedingY' parameter represents the summer breeding season, and 'SexM' as male birds and 'SexU' as birds of undetermined sex.

| | AICc | df | Variable | Estimate | SE | *t*-value | $x^2$ | df | *p*-value |
|---|---|---|---|---|---|---|---|---|---|
| Full Model | 184.4 | 9 | Intercept | 14.106 | 0.290 | 48.66 | | | |
| | | | Elevation | -0.161 | 0.140 | -1.15 | 1.321 | 1 | 0.250 |
| | | | SexM | -0.155 | 0.364 | -0.43 | 2.308 | 2 | 0.315 |
| | | | SexU | -0.910 | 0.597 | -1.52 | | | |
| | | | BreedingY | -1.055 | 0.601 | -1.77 | 0.515 | 1 | 0.472 |
| | | | SexM* BreedingY | 1.085 | 0.688 | 1.58 | 2.600 | 2 | 0.273 |
| | | | SexU* BreedingY | 1.225 | 1.313 | 0.93 | | | |
| Null Model | 168.3 (Δ0) | 3 | Intercept | 13.859 | 0.143 | 97.05 | | | |
| | 170.3 (Δ2.01) | 4 | Intercept | 13.953 | 0.169 | 82.46 | | | |
| | | | BreedingY | -0.319 | 0.300 | -1.06 | 1.127 | 1 | 0.288 |
| | 171 (Δ2.74) | 4 | Intercept | 13.863 | 0.143 | 97.19 | | | |
| | | | Elevation | -0.192 | 0.139 | -1.37 | 1.898 | 1 | 0.168 |
| | 171 (Δ2.78) | 5 | Intercept | 13.853 | 0.255 | 54.35 | | | |
| | | | SexM | 0.106 | 0.315 | 0.34 | 2.026 | 2 | 0.363 |
| | | | SexU | -0.613 | 0.535 | -1.15 | | | |
| | 172.7 (Δ4.39) | 8 | Intercept | 14.118 | 0.289 | 48.84 | | | |
| | | | SexU | -0.134 | 0.366 | -0.37 | 2.347 | 2 | 0.3101 |
| | | | SexM | -0.910 | 0.596 | -1.53 | | | |
| | | | BreedingY | -1.126 | 0.596 | -1.89 | 1.171 | 1 | 0.279 |
| | | | SexM* BreedingY | 1.050 | 0.695 | 1.51 | 2.465 | 2 | 0.292 |
| | | | SexU* BreedingY | 1.285 | 1.309 | 0.98 | | | |
| | 172.9 (Δ4.67) | 6 | Intercept | 13.939 | 0.263 | 53.04 | | | |
| | | | SexM | 0.144 | 0.315 | 0.46 | 2.353 | 2 | 0.308 |
| | | | SexU | -0.626 | 0.531 | -1.18 | | | |
| | | | BreedingY | -0.366 | 0.302 | -1.21 | 1.471 | 1 | 0.225 |
| | 173.7 (Δ5.41) | 5 | Intercept | 13.932 | 0.169 | 82.52 | | | |
| | | | Elevation | -0.156 | 0.143 | -1.09 | 1.196 | 1 | 0.274 |
| | | | BreedingY | -0.237 | 0.297 | -0.80 | 0.636 | 1 | 0.425 |
| | 173.9 (Δ5.62) | 6 | Intercept | 13.859 | 0.254 | 54.52 | | | |
| | | | Elevation | -0.197 | 0.139 | -1.42 | 2.009 | 1 | 0.156 |
| | | | SexM | 0.105 | 0.314 | 0.33 | 2.128 | 2 | 0.345 |
| | | | SexU | -0.631 | 0.533 | -1.18 | | | |

**Table S11** Tukey post-hoc comparison tests of blood haemoglobin concentrations ($H_b$ gd/L) between species per season from a linear mixed effect model of $H_b$ as a function of species. Significant comparisons are emboldened and acronyms follow those in main text (ruddy-capped nightingale-thrush *Catharus frantzii*, RCNT; black-headed nightingale-thrush *Catharus mexicanus*, BHNT; chestnut-capped brushfinch *Arremon brunneinucha*, CCBF; grey-breasted wood wren *Henicorhina leucophrys*, GBWW; common bush tanager *Chlorospingus flavopectus*, COBT).

| Species comparison | Estimate | SE | z-value | *p*-value |
|---|---|---|---|---|
| **Summer (breeding season)** | | | | |
| RCNT - CCBF | 1.714 | 0.424 | 4.042 | **<0.001** |
| BHNT - CCBF | 0.057 | 0.309 | 0.185 | 1.000 |
| COBT - CCBF | -0.214 | 0.375 | -0.564 | 0.979 |
| GBWW - CCBF | -0.770 | 0.419 | -1.838 | 0.341 |
| BHNT - RCNT | -1.657 | 0.348 | -4.768 | **<0.001** |
| COBT - RCNT | -1.926 | 0.408 | -4.724 | **<0.001** |
| GBWW - RCNT | -2.484 | 0.448 | -5.542 | **<0.001** |
| COBT - BHNT | -0.269 | 0.286 | -0.941 | 0.875 |
| GBWW - BHNT | -0.827 | 0.344 | -2.424 | 0.104 |
| GBWW - COBT | -0.558 | 0.401 | -1.387 | 0.625 |
| **Winter (non-breeding season)** | | | | |
| BHNT - CCBF | 0.176 | 0.351 | 0.502 | 0.955 |
| COBT - CCBF | 0.399 | 0.315 | 1.270 | 0.559 |
| GBWW - CCBF | -0.224 | 0.761 | -0.294 | 0.990 |
| COBT - BHNT | 0.224 | 0.278 | 0.804 | 0.841 |
| GBWW - BHNT | -0.400 | 0.747 | -0.536 | 0.946 |
| GBWW - COBT | -0.624 | 0.731 | -0.854 | 0.816 |
